# Multiple paralogues and recombination mechanisms drive the high incidence of 22q11.2 Deletion Syndrome

**DOI:** 10.1101/2024.03.14.585046

**Authors:** Lisanne Vervoort, Nicolas Dierckxsens, Marta Sousa Santos, Senne Meynants, Erika Souche, Ruben Cools, Tracy Heung, Koen Devriendt, Hilde Peeters, Donna M. McDonald-McGinn, Ann Swillen, Jeroen Breckpot, Beverly S. Emanuel, Hilde Van Esch, Anne S. Bassett, Joris R. Vermeesch

## Abstract

The 22q11.2 deletion syndrome (22q11.2DS) is the most common microdeletion disorder. Why the incidence of 22q11.2DS is much greater than that of other genomic disorders remains unknown. Short read sequencing cannot resolve the complex segmental duplicon structure to provide direct confirmation of the hypothesis that the rearrangements are caused by non-allelic homologous recombination between the low copy repeats on chromosome 22 (LCR22s). To enable haplotype-specific assembly and rearrangement mapping in LCR22 regions, we combined fiber-FISH optical mapping with whole genome (ultra-)long read sequencing or rearrangement-specific long-range PCR on 24 duos (22q11.2DS patient and parent-of-origin) comprising several different LCR22-mediated rearrangements. Unexpectedly, we demonstrate that not only different paralogous segmental duplicon but also palindromic AT-rich repeats (PATRR) are driving 22q11.2 rearrangements. In addition, we show the existence of two different inversion polymorphisms preceding rearrangement, and somatic mosaicism. The existence of different recombination sites and mechanisms in paralogues and PATRRs which are copy number expanding in the human population are a likely explanation for the high 22q11.2DS incidence.

## Introduction

Low copy repeats (LCRs), also referred to as segmental duplications, constitute 6.6% of the human genome (Nurk et al. 2022) and played an important role during human evolution and adaptation (Dennis and Eichler 2016). They are defined as DNA segments with a length of at least 1 kb that share >90% of sequence identity (Bailey et al. 2001, 2002). Their high sequence homology is known to be a driver of non-allelic homologous recombination (NAHR), caused by meiotic misalignment of homologous chromosomes or sister chromatids, resulting in reciprocal deletions, duplications, and inversions (Inoue and Lupski 2002; Porubsky et al. 2022). These recurrent genomic rearrangements often cause genomic disorders, forming collectively an important cause of disabling diseases in the general population (Angelis et al. 2015).

The 22q11.2 deletion syndrome (22q11.2DS, MIM 188400) is the most common microdeletion in humans with an estimated incidence of 1 in 2148 live births (Blagojevic et al. 2021). 22q11.2DS has a heterogeneous presentation including multiple congenital and later-onset features, such as cardiac, palatal, metabolic, cognitive, and neuropsychiatric abnormalities (McDonald-McGinn et al. 2015). The presence and severity of the associated clinical expression is variable but the causes of this high variability remain largely unexplained (McDonald-McGinn et al. 2015). Extensive investigation of the recombination locus is complex due to the presence of eight LCRs on the chromosome, commonly named LCR22-A until -H (Shaikh et al. 2000). Rearrangements between different segments occur (Campbell et al. 2018; McDonald-McGinn et al. 2015), but deletions of the 3Mb region extending from LCR22-A to -D represent the main cause (85%) of the 22q11.2DS (Campbell et al. 2018). CNVs distal to LCR22-D represent a separate condition (MIM 611867).

Why the incidence of the 22q11.2DS is an order of magnitude higher than any other genomic disorder remains an enigma. Although it is assumed that the rearrangements are caused by NAHR between the LCR22s, direct confirmation of this hypothesis is lacking. This is because for a long time, a gapless reference of the LCR22s was missing. The gaps were a consequence of a complex intricate segmental duplicon structure which could not be resolved by short read sequencing techniques and the lack of bioinformatic pipelines to create haplotype-resolved LCR22 assemblies (Vollger et al. 2019). Several attempts to map the rearrangements have provided anecdotal results: using long-range PCR and sequencing of two distal 22q11.2 deletions, the Breakpoint Cluster Region (BCR) module was suggested to be the NAHR site in the distal LCR22-D/E and LCR22-E/F deletions (Shaikh et al. 2007). By mapping shared and paralogous sequence polymorphisms of LCR22-A and -D by sequencing bacterial artificial chromosomes (BACs), this same BCR module was suggested to also drive the NAHR in a single LCR22-A/D deletion patient (Guo et al. 2016).

With the help of optical mapping techniques, Demaerel et al. (2019) were able to uncover the complex repeat structure of the LCR22s in the human population. They are characterized by a high variability in size (200kb – 3Mb) and structural organization of the LCR22-A haplotype(Demaerel et al. 2019; Pastor et al. 2020). Most recently, the telomere-to-telomere (T2T) consortium released the first gapless fully sequenced haploid genome, including complete LCR22s (Nurk et al. 2022). Although the sequence represents an existing LCR22-A haplotype, the allele is rather short and some specific duplicons, present in part of the human population, are missing. Hence, this initial T2T reference cannot be used as an accurate representation in LCR22-mediated NAHR research.

NAHR in the BCR modules cannot explain the high incidence of the 22q11.2DS, raising the question if additional mechanisms are responsible for the LCR22-mediated rearrangements. To enable LCR22 haplotype-specific assembly and rearrangement mapping in those regions, we combined optical mapping with whole genome (ultra-)long read sequencing and/or rearrangement-specific long-range PCR. A combination of these methods was applied on 24 duos (patient and parent-of-origin, **Supplemental_Table_S1**) and one index patient with five different LCR22-mediated rearrangements to map the crossover sites. Unexpectedly, we demonstrate that not only highly identical LCR22 segments, but also palindromic AT-rich repeat (PATRR) instability, are driving 22q11.2 rearrangements.

## Results

### Fiber-FISH identification of LCR22-specific recombination clusters

To map NAHR sites, we first generated *de novo* assembly fiber-FISH or optical maps of all LCR22-A, -B, -C and -D alleles in parental and patient genomes (Demaerel et al. 2019) (**Fig. 1**). Fiber-FISH provides haplotype-aware *de novo* assembly at subunit resolution with long-range structural information of the targeted LCR22 loci. In addition, the loci can be assembled without a priori structural knowledge of the LCR22s, tackling assembly problems of regions associated with incorrect reference genome representation and extreme structural variation.

**Fig. 1:**
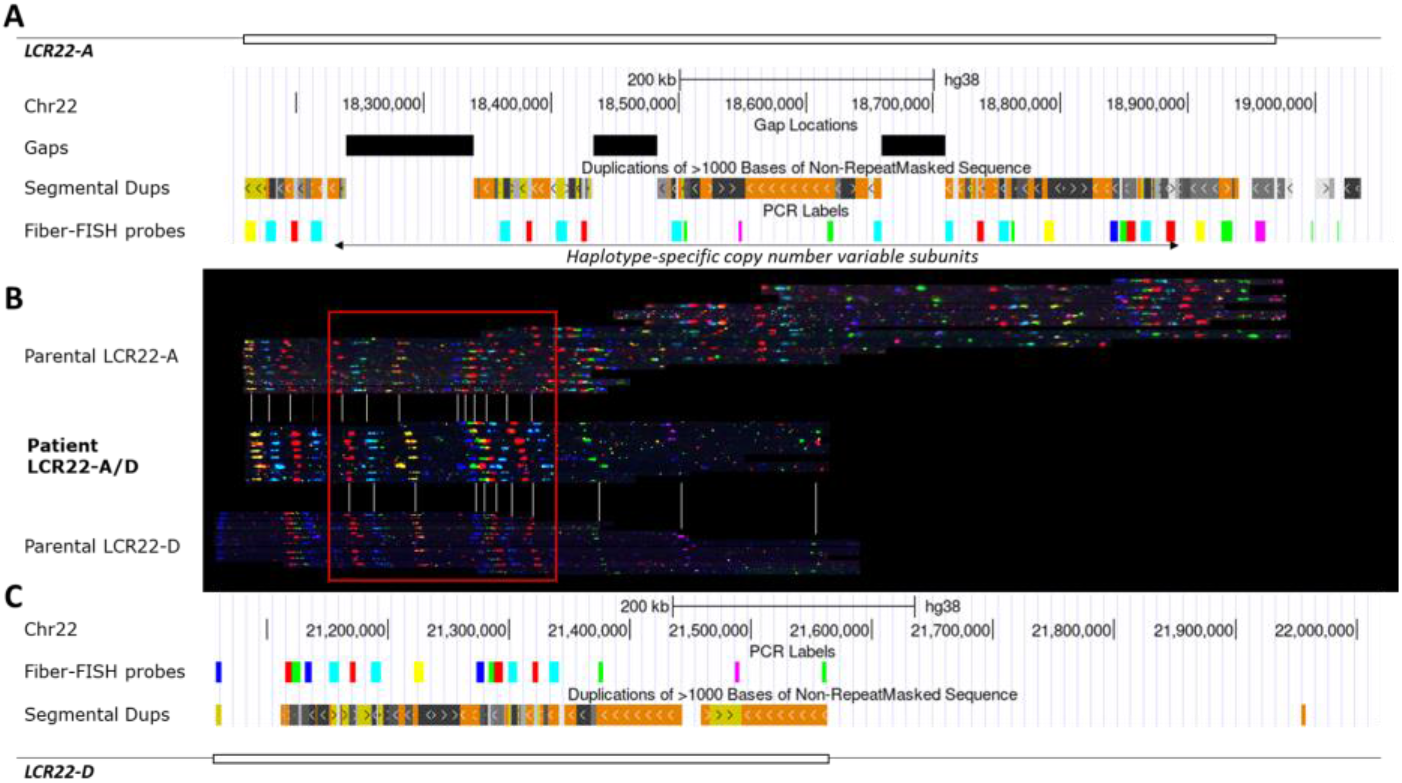
Fiber-FISH analysis of a LCR22-mediated (A-D) rearrangement. (A) UCSC Genome browser screenshot of hg38 with tracks for gaps, segmental duplications and blat sequences for fiber-FISH probes in LCR22-A. (B) LCR22-A/D rearrangement with *de novo* assembled parental LCR22-A, -D (parent-of-origin), and 22q11.2DS patient rearrangement alleles. The parental LCR22-A haplotype is not identical to the reference allele in A, due to presence of gaps in the reference and haplotype variability observed in the human population. The sequence in the red box is shared between three alleles and therefore considered to be the locus where recombination had taken place. Gray lines indicate similarities between the alleles of the patient and the parent-of-origin. (C) UCSC Genome browser screenshot of hg38 with tracks for fiber-FISH probes and segmental dups in LCR22-D.

Crossover regions were determined in 24 families, by comparison of the rearrangement alleles in patient and parent-of-origin **(Supplemental_Fig_S1, Supplemental_Fig_S2 and Supplemental_Fig_S3)**. Our sample collection included five different LCR22-mediated recombinations: 15 LCR22-A/D deletions, five LCR22-A/B deletions (four patient-parent duos and one index patient), three LCR22-A/C deletions, one LCR22-B/D deletion and, one LCR22-C/D deletion **(Table S1)**. Rearrangements occurred for nine families on the maternal allele and for 15 families on the paternal allele. Fiber-FISH patterns of the patient and the parent-of-origin, in whom the rearrangement occurred, were analyzed. All parental alleles showed normal heterozygous LCR22-A, homozygous LCR22-B and -C, and hetero- or homozygous LCR22-D patterns, as described in Demaerel et al. (2019) and Pastor et al. (2020). In addition, an inversion between LCR22-A and -D was observed in the parent-of-origin of one family (BD001) **(Fig. 4A, Supplemental_Fig_S1)**.

Based on *in silico* predictions, several of the segmental duplicons identified could act as NAHR substrate, and these duplicon NAHR sites cluster in a subset (**Table 1, Supplemental_Figure_S1**). We chose to use reference genome hg38 for comparison and visualization of the LCR22s. For nested deletions involving LCR22-C, the rearrangements cluster in a 10kb region (Chr22: 20,688,715-20,698,995), which is copy number variable in LCR22-A and -D **(Table 1)**, from now on named RL-C (recombination locus LCR22-C). LCR22-B-mediated nested deletions can be subcategorized into two groups: (I) two out of five crossovers occurred in a 20kb unit (Chr22: 20,324,573-20,344,531; RL-B1), and (II) in three out of five the region was refined to an 8kb locus (Chr22: 20,331,987-20,339,583; RL-B2) within this 20kb unit. The 20kb unit is copy number variable in both LCR22-A and LCR22-D (Demaerel et al. 2019). Due to the size and the structural variation, rearrangements involving both LCR22-A and -D are the most complex to analyze. In 13 out of 15 families, recombination occurs within a 160kb locus (RL-AD1) whereas in the other three families, the recombination was restricted to a 20kb (RL-AD2) interval within RL-AD1. All LCR22 recombination loci identified are part of the larger RL-AD1 locus (**Table 1**).

**Table 1:** 22q11.2 LCR rearrangement loci at fiber-FISH resolution. The chromosomal locus corresponds to the genomic location in reference genome hg38. CNV in superscript indicates the presence of copy number variability of the recombination locus in the LCR22 (-A and -D), based on Demaerel et al. (2019) and Pastor et al. (2020).

| Deletion type | # of samples | Recombination locus | Locus in proximal LCR22 | Locus in distal LCR22 |
| --- | --- | --- | --- | --- |
| LCR22-A/D | 12 | RL-AD1 (160kb) | chr22:18,729,876-18,891,257<sup>cnv</sup> | chr22:21,162,204-21,325,236<sup>cnv</sup> |
|  | 2 | RL-AD2<br>(20kb) | chr22:18,838,944-18,859,182<sup>cnv</sup> | chr22:21,115,387-21,137,124<sup>cnv</sup> |
| **LCR22-A/B** | 2 | RL-B1<br>(20kb) | chr22:18,196,206-18,218,744<sup>cnv</sup> | chr22:20,324,573-20,344,531 |
|  | 3 | RL-B2<br>(8kb) | chr22:18,414,455-18,422,770<sup>cnv</sup> | chr22:20,331,987-20,339,583 |
| **LCR22-A/C** | 3 | RL-C<br>(10kb) | chr22:18,845,620-18,855,844<sup>cnv</sup> | chr22:20,688,715-20,698,995 |
| **LCR22-B/D** | 1 | RL-B1<br>(20kb) | chr22:20,324,573-20,344,531 | chr22:21,152,820-21,173,481<sup>cnv</sup> |
| **LCR22-C/D** | 1 | RL-C<br>(10kb) | chr22:20,688,715-20,698,995 | chr22:21,279,982-21,291,168<sup>cnv</sup> |

### Long-read sequencing toolkit to map *22q11*.*2* recombinations

Since fiber-FISH resolution is limited, we leveraged long-read sequencing and optimized a *de novo* assembler algorithm to scrutinize these crossovers in more detail (**Fig. 2**). Long read sequencing was performed in ten of the 24 families. Ultra-long read sequencing (ULK, **Fig. 2B**) was performed in eight. N50 values were over 50kb (maximum of 132kb) with an output range of 4-111Gb. To increase coverage, the ULK data were complemented by standard-long read sequencing (SLK, **Fig. 2B**) and/or High Duplex (HD) sequencing in three families. For one family, only standard-long sequencing was performed. SLK and HD sequencing resulted in 36-115Gb and 96Gb of sequencing data respectively, with an N50 value above 21kb (**Supplemental_Table_S2**). This allowed the identification of the rearrangement breakpoint in all nine families (**Supplemental_Table_S1**). In addition, we were able to infer the crossover site by long-range PCR and subsequent long-read sequencing in one index patient (AB004, **Fig. 2B**) with a LCR22-A/B deletion.

**Fig. 2:**
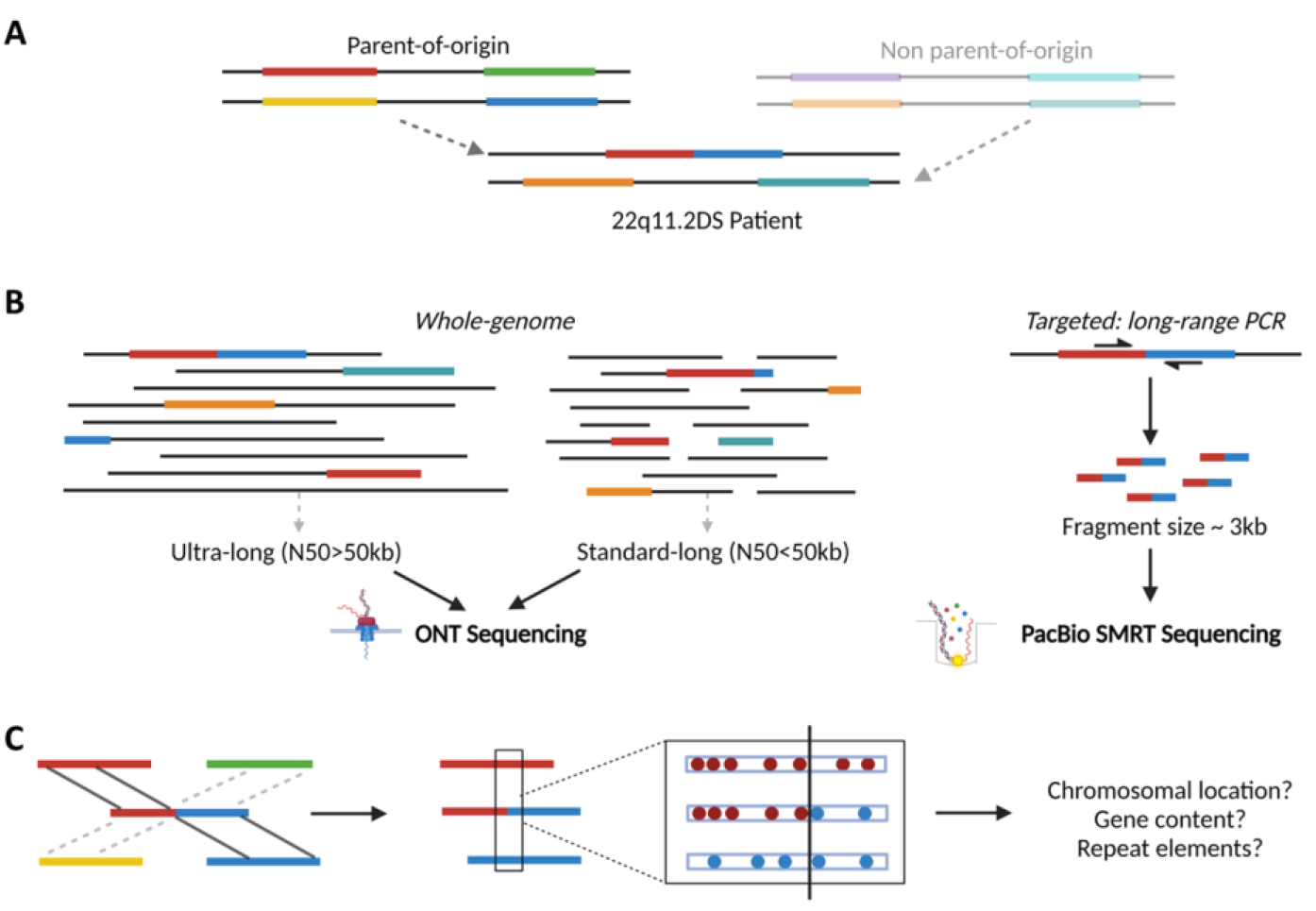
Study design and data analysis of 22q11.2DS recombination loci. **(A)** LCR22 composition of a family trio. The parent-of-origin is the parent in whom the recombination occurred. The red-blue composition in the patient represents the recombined LCR22 structure. In this example, an interchromosomal recombination is presented, although intrachromosomal recombinations are possible as well. **(B)** The patient and the parent-of-origin were sequenced using one or a combination of whole-genome ultra-long and/or standard-long Oxford Nanopore Technologies (ONT) sequencing. In one patient (AB004) a targeted breakpoint-specific long-range PCR was designed in combination with PacBio single-molecule real-time (SMRT) sequencing of the fragment. **(C)** Following *de novo* assembly of the different proximal and distal LCR22 alleles in the patient and parent-of-origin, the recombination-involved parental haplotypes (proximal and distal) were identified via SNP comparison, followed by a delineation of the breakpoint locus by LCR22 proximal-specific (red) and distal-specific (blue) SNPs. These SNPs are shared between the patient and the parental proximal or distal LCR22, respectively, but not with the other (proximal or distal) LCR22 involved in the CNV. The recombination locus was scrutinized or its precise coordinates, elucidating the genes and repeat elements involved.

Existing long read assembly algorithms are graph-based and are therefore designed to assemble the complete genome. Without additional short-sequencing reads for trio binning or Hi-C data, phasing remains problematic. The recent efforts of the T2T consortium have proven that assembling even a relatively short LCR22-A is not a trivial task. From the different assemblies of their recently published haplotype-resolved human genome, only the ones that combine both HiFi and ONT reads at sequencing depths several magnitudes higher than regular datasets were able to resolve the LCR22-A (Rautiainen et al. 2023). Since existing assembly algorithms were not able to resolve any of the LCR22 regions, we developed a new targeted haplotype-aware assembler. Considering that we are interested in assembling relatively short regions as accurately as possible, we opted for a seed-and-extend method that assembles both haplotypes in parallel, starting from a given seed sequence. The algorithm keeps track of the variation between the haplotypes and it looks for mismatch patterns between the reads and each haplotype assembly to select the correct path during each iteration **(Fig. 2C)**.

### Variability of the crossover site at nucleotide resolution

We leveraged these long-read sequencing approaches to resolve the recombination alleles for the rearrangements occurring in nine of the 24 families (**Fig. 3, Table 2**, and **Supplemental_Fig_S4**). As opposed to the larger, apparently uniform, blocks identified by optical mapping, the breakpoint loci identified by this approach were scattered within these blocks (**Fig. 3**) and located in a variety of gene loci and involving various repeat elements (**Table 2, Supplemental_Fig_S4**).

**Table 2:**
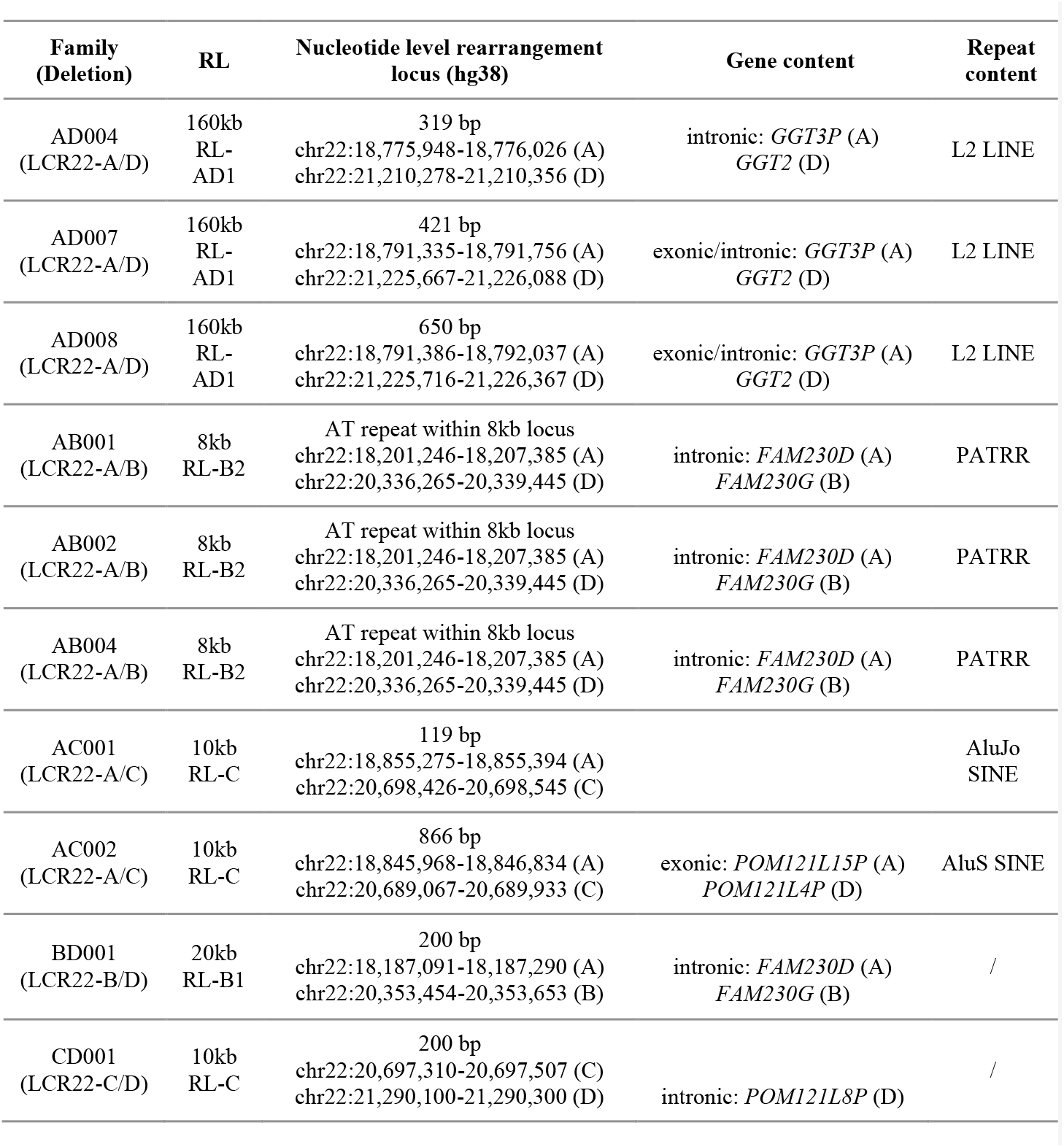
22q11.2 LCR rearrangement loci (RL) at nucleotide resolution. First column shows the family identifier, based on **Supplementary_Table_1**, and diagnosed deletion type. The fiber-FISH based (**Table 1**) as well as the length of the RL is presented in the second column. The third column provides the position of the last proximal-specific SNP and the first distal-specific SNP and the length of the locus. The Exact position could not be mapped for the three PATRR-mediated LCR22-A/B deletions due to presence of several PATRRs in the reference genome. Columns four and five show the gene and repeat content, respectively. Repeat elements and gene exons/introns can be fully or partly covered in the recombination locus. More information on the exact composition of the recombination locus can be found in **Supplementary_Fig_3**. In family BD001, the deletion is diagnosed as LCR22-B/D, however, the recombination occurred between LCR22-A and -B (corresponding to chromosomal loci in the third column), as explained in **Figure 4C**.

**Fig. 3:**
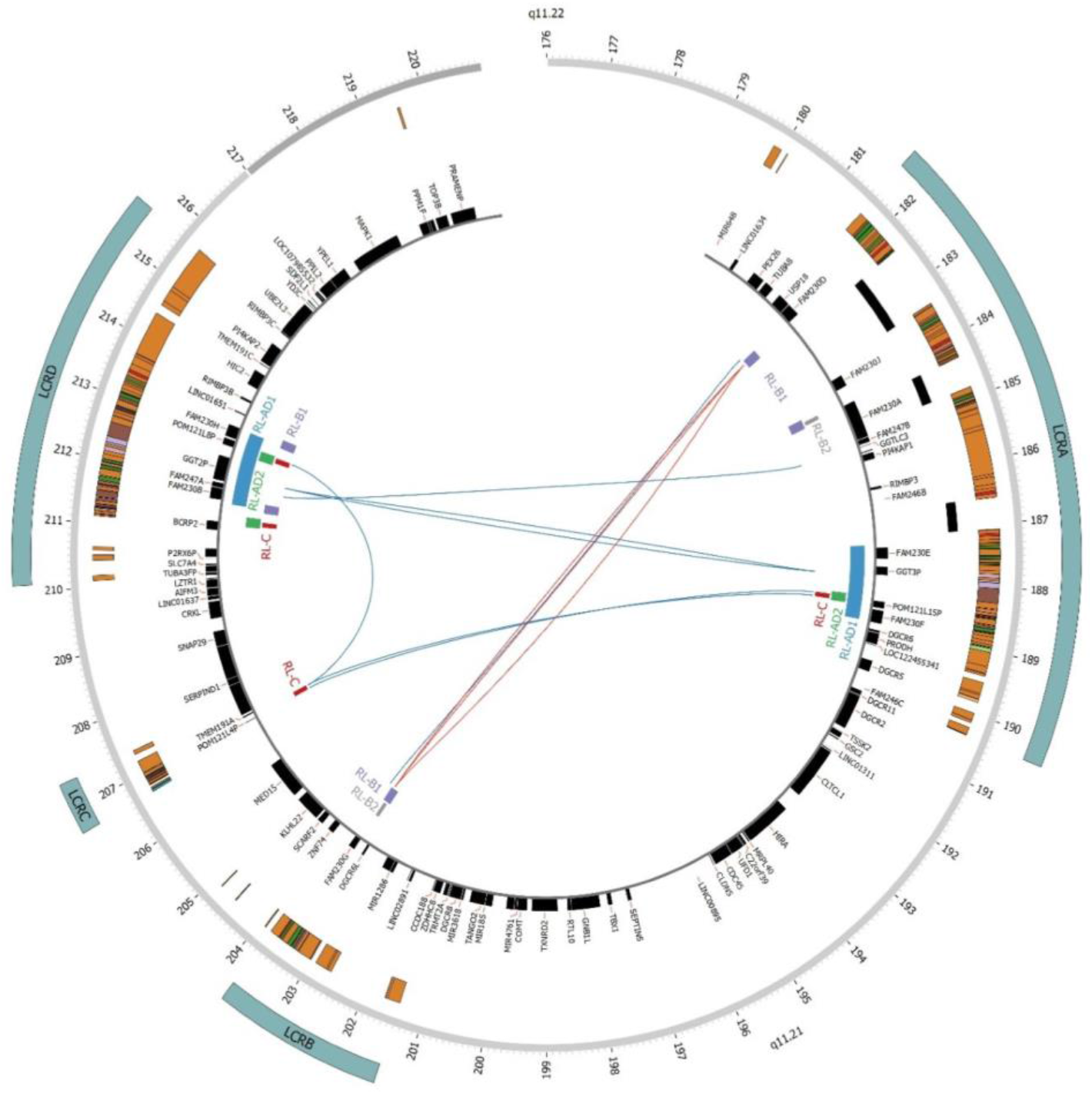
Schematic visualization of 22q11.2 recombination breakpoints. The graph represents the 22q11.2 locus, its segmental duplications and known genes. Lines between LCR22s connect the proximal and distal breakpoint of a family-specific recombination. Solid red lines represent PATRR-mediated recombinations, and solid blue lines represent NAHR-mediated recombinations. The position of genes located in the involved recombination sequences are shown via blue or green vertical dotted lines.

In three out of the five LCR22-A/B deletions, the fiber-FISH pattern showed a clear transition from LCR22-A to -B in absence of large segmental duplicon structures of >20kb where NAHR could have taken place **(Supplemental_Fig_1, Table 1)**. Using ultra-long read sequencing (patients AB001 and AB002) and single-molecule real-time long-read sequencing (patient AB004), we revealed that the crossover occurred within a palindromic AT-rich repeat (PATRR). In addition, in family AB001, a 50bp LINE element, which was not present on either parental allele, was inserted at the recombination site between two PATRRs. The presence of a LINE insertion and the involvement of the PATRRs provides evidence for non-homologous end-joining (NHEJ) as a cause of 22q11.2 deletion.

### Complex recombination patterns mediated by LCR22 inversions

In a family (BD001) with a *de novo* LCR22-B/D deletion, the fiber-FISH LCR22 structures showed (I) a normal LCR22-A, -B, -C, and -D, (II) an LCR22 block with the proximal start of LCR22-A combined with the proximal part of LCR22-B in an inverted orientation (LCR22-A/Binv), and (III) the distal end of LCR22-A in an inverted orientation coupled to the distal end of LCR22-D (LCR22-Ainv/D) **(Fig. 4A, Supplemental_Fig_S1)**. In the parent-of-origin, the normal LCR22-A, -B, -C, and -D alleles were observed, as well as an allele indicative of an LCR22-A/D inversion **(Fig. 4A, Supplemental_Fig_S1)**. Interphase-FISH using probes proximal and distal from LCR22-A and proximal from LCR22-B show the presence of an inversion in 100% of the cells of both the patient and the parent-of-origin **(Fig. 4B, Supplemental_Table_S3)**. Indeed, the presence of a parental LCR22-A/D inversion in combination with a NAHR recombination between LCR22-A and -B does explain the observed fiber-FISH structures in the patient. Using long-read sequencing, this recombination was shown to be intronic in the FAM230 paralogue in both LCR22-A and -B. As a consequence, the locus between LCR22-B and -D is deleted. In a family (AB002) with a *de novo* LCR22-A/B deletion, a mosaic LCR22-A/B deletion was identified via low-pass sequencing, and validated by arrayCGH, SNP array, and dual color interphase-FISH (TUPLE1/Arsa). Fiber-FISH uncovered the presence of three different alleles in the patient: (I) a normal LCR22-A and -B haplotype, (II) the LCR22-A/B deletion haplotype, and (III) a haplotype carrying an inversion between LCR22-A and -B **(Fig. 4C, Supplemental_Fig_S1)**. The fiber-FISH patterns in the parent-of-origin showed two normal alleles **(Fig. 4C, Supplemental_Fig_S1)**. In addition, interphase-FISH validated the presence of the three haplotypes and uncovered that each cell carried a wild type 22q11.2 locus and either the LCR22-A/B deletion or inversion **(Fig. 4D, Supplemental_Table_S3)**. We hypothesize the LCR22-A/B deletion was created from the LCR22-A/B inversion allele in an early stage during embryogenesis. Based on whole-genome ultra-long read sequencing data of the family, the deletion crossover was pinpointed in a PATRR of LCR22-A and -B of the inversion allele. Similarly, these PATRRs seem to be responsible for the creation of the inversion allele as well **(Fig. 4C)**. The most parsimonious explanation is that two consecutive PATRR-mediated events created an LCR22-A/B inversion and a deletion allele in the patient from family AB002 **(Fig. 4C)**.

**Fig. 4:**
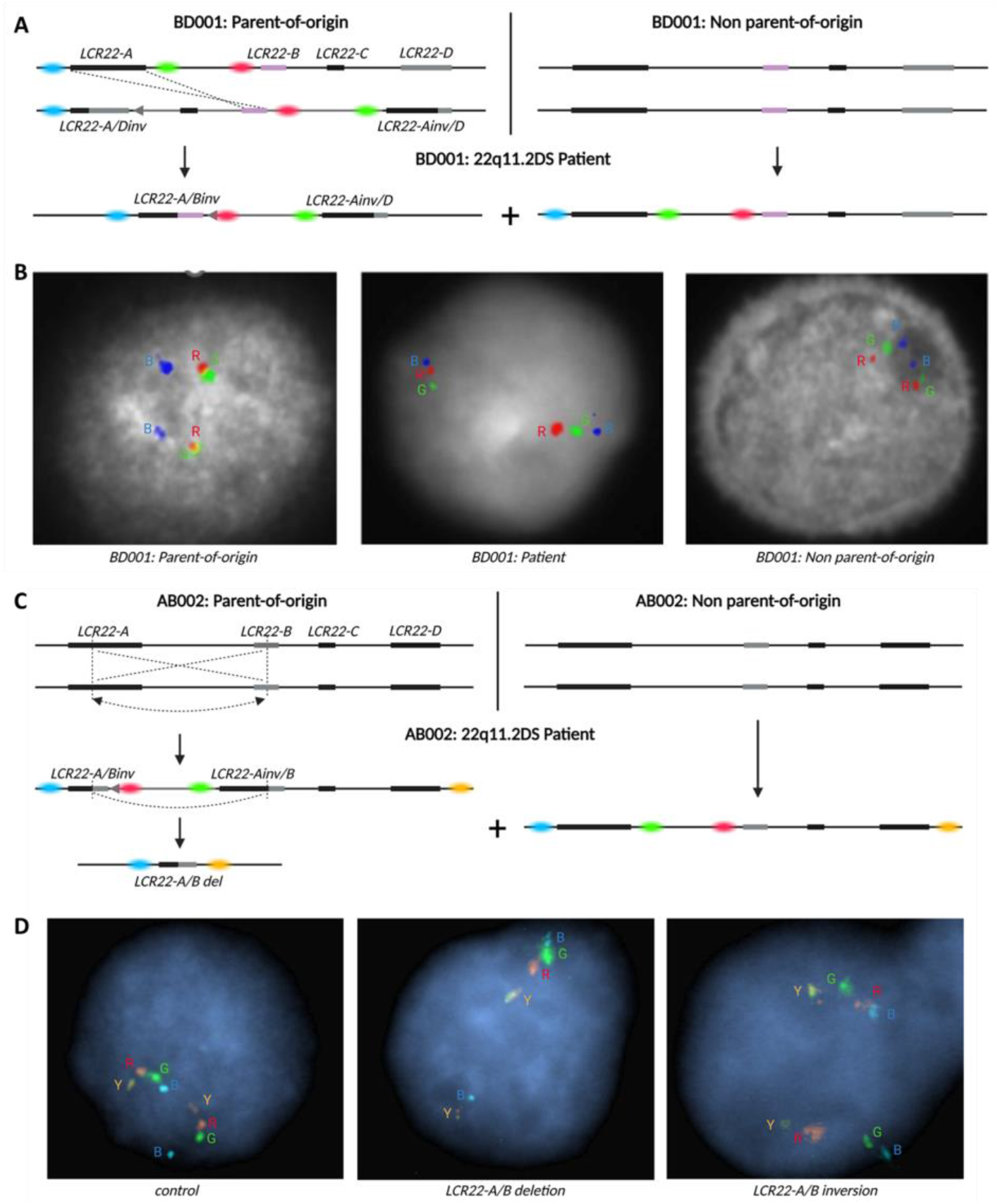
Inversion-associated rearrangements. **(A)** Schematic representation of the rearrangement of family BD001: the parent-of-origin (left) carries a LCR22-A/D inversion, which is recombined between LCR22-A and - B (dotted line) causing the presence of a deletion allele in the patient, missing the sequence from LCR22 -B until -D. In addition, the patient inherited one of the wild type alleles of the other parent (right). Colored dots represent the probes used in the interphase-FISH. **(B)** Interphase-FISH results using color-labeled probes CH17-320A22 (proximal LCR22-A, blue, B), CH17-222C16 (distal LCR22-A, green, G), and CH17-389E17 (proximal LCR22-B, red, R) in the parent-of-origin (left), patient BD001 (middle), and the other parent (right). **(C)** Schematic representation of the rearrangement of family AB002: both parents carry two normal LCR22 haplotypes (LCR22-A until -D). The patient carries a wild type allele (right) and an LCR22-A/B inversion allele is observed (inverted gray arrow), created by the recombination (dotted line and arrow) of the LCR22-A and -B allele from the parent-of-origin. We hypothesize the deletion is created in a consecutive manner by a new recombination between these blocks. Colored dots represent the probes used in the interphase-FISH. **(D)** Interphase-FISH results using color-labeled probes CH17-320A22 (proximal LCR22-A, blue, B), CH17-222C16 (distal LCR22-A, green, G), CH17-389E17 (proximal LCR22-B, red, R), and RP11-354K13 (distal LCR22-D, yellow, Y) in a control individual (left), deletion-carrying cell of patient AB002 (middle), and inversion-carrying cell of patient AB002 (right).

## Discussion

Due to the limitations of short- and standard long-read sequencing methods and the absence of an accurate reference genome, the exact positions of the recombinations of the 22q11.2DS remained uncharted. As a consequence, the mechanisms involved in the recombination and the reason for the high incidence of 22q11.2 rearrangements remained unknown. In this study, we mapped the recombination sites in 24 families with *de novo* 22q11.2 CNVs and were able to map the recombination sites at the nucleotide resolution in ten families using long-read sequencing approaches. We demonstrate that different paralogues in the LCR22s can drive recombination. Most paralogous are copy variable in the population. Interestingly, we reveal that different rearrangements occur in the PATRR loci within the LCR22s, suggesting that not only NAHR but also breakage-mediated repair mechanisms occur and likely contribute to the high incidence of the syndrome.

PATRRs are palindromic sequences that create genomic instability via the formation of single-stranded hairpin or double-stranded cruciform secondary structures. Those cruciforms are sensitive to the generation of double-strand breaks, which are repaired via NHEJ (Kato et al. 2012). The LCR22-B PATRR is known to drive recurrent 22q11.2 translocations (Kurahashi et al. 2006), but has not been reported to be involved in 22q11.2 deletions. Here, we show involvement of PATRR sequence located within LCR22-A and -B to create LCR22-A/B deletions. It is known that PATRR size polymorphisms influence the rearrangement frequency of *de novo* t(11;22) translocations (Kato et al. 2006; Tong et al. 2010). For example, larger and symmetric PATRRs on chromosomes 11 and 22 are more prone to t(11;22) translocations (Kato et al. 2006; Tong et al. 2010). Interestingly, we uncovered structural polymorphisms in PATRR tandem repeats, with copy numbers ranging between 0 and 13 and variability in the AT repeat lengths in the four investigated individuals (patient and parent in families AB001 and AB002 with *de novo* LCR22-A/B deletions, data not shown). It seems likely that this repeat variability could affect rearrangement frequency. Also, Correll-Tash et al. (2021) showed that secondary structure formation and double-strand breaks can occur both during meiosis and mitosis. Here, we identified an individual with a possible meiotic, followed by a mitotic progression of PATRR-driven rearrangements (AB002, **Fig. 4C**). Alternatively, both the inversion and the deletion could have occurred post-zygotically, but this possibility appears less likely given the absence of a non-rearranged LCR22 haplotype inherited from the parent-of-origin.

Different segmental duplicons contribute to NAHR events at the 22q11.2 locus (**Table 2**). Hence, various recombination loci may be present in the shared modules between the two involved LCR22s (proximal and distal) and these create variability of the crossover locus. The identification of multiple subunits driving NAHR is also observed in other genomic disorders, for example neurofibromatosis type I deletions (Summerer et al. 2018), and Sotos syndrome 5q35 CNVs (Visser et al. 2005a, 2005b). In 22q11.2DS, in addition, many of the NAHR and PATRR sites appear to be copy number variable. It thus seems likely that expanded haplotypes may be more likely to predispose to rearrangement events. This hypothesis can now be tested by analyzing LCR22s in a larger (22q11.2DS) population.

Among the families studied, we identified two individuals with LCR22-mediated inversions, one a parent-of-origin with a LCR22-A/D inversion that led to a LCR22-B/D deletion offspring, and the other a patient with a mosaic genotype that included a LCR22-A/B inversion. Previous studies had failed to identify such inversions in 22q11.2DS (Gebhardt et al. 2003; Vergés et al. 2017). However, availability of a gapless telomere-to-telomere assembly of the human genome (T2T-CHM13) has enabled a more complete analysis of genomic variation (Nurk et al. 2022). Using long read sequencing and Strand-seq data of 52 samples from the 1000 Genomes project and mapping these against the T2T-CHM13 reference has provided a more complete view of the genome-wide inversion polymorphism (Porubsky et al. 2022; Hanlon et al. 2022; Porubsky et al. 2023). Although not directly related to disease, LCR-mediated inversion polymorphisms drive NAHR, thus can lead to genomic disorders (Shaw and Lupski 2004). Examples of this phenomenon can be found in the high incidence of inversion-carrying parents-of-origin in Williams-Beuren syndrome (Osborne et al. 2001), Angelman syndrome, Sotos syndrome, 8p23.1 microdeletion, and 15q23 and 15q24 microdeletion syndromes (Puig et al. 2015). In some cases, these disease-predisposing inversion polymorphisms can be linked to phenotypic consequences (Boettger et al. 2012; Steinberg et al. 2012). To address the incidence and eventual phenotypic consequences in 22q11.2DS, it will be critical to survey many more human genomes and to sequence and resolve the large complex LCR22s flanking the inversion polymorphisms, along with deep phenotyping data.

We observed one individual to be mosaic, with 50% white blood cells to have a normal oriented LCR22 haplotype, 25% a LCR22-A/B inversion haplotype, and 25% a LCR22-A/B deletion. Mosaicism of the 22q11.2 deletion is rare, with few cases previously reported (Consevage et al. 1996; Halder et al. 2008; Chen et al. 2019; Patel et al. 2006; Chen et al. 2004). These reports were based on standard interphase or metaphase FISH testing, and thus no recombination sites were mapped. Interestingly, one case described 22q11.2 deletion mosaicism in a miscarried fetus (85% of cells) as well as in the mother (11% of cells), suggesting an increased recombination susceptibility for the specific chromosome involved (Patel et al. 2006). It will be of interest to map the LCR22 haplotypes of other individuals with 22q11.2DS mosaicism. We hypothesize inversions and PATRRs to be drivers of postzygotic rearrangements.

In conclusion, we uncovered crossovers occur in different paralogues within the LCR22s and both NAHR and NHEJ can cause 22q11.2DS. However, the large and complex LCR22s -A and -D remain challenging to sequence which resulted in only a limited number of recombinations resolved, despite being the most frequent. With improved and cheaper long read sequencing technologies, it becomes feasible to *de novo* assemble the ultra-long and complex LCR22s which, in turn, will enable to saturate the landscape of 22q11.2 rearrangements. Since haplotype variability of segmental duplicons has been shown to affect rearrangement predisposition for other genomic disorders (Steinberg et al. 2012), exploration of the size, numbers and orientation of paralogue variability can potentially uncover predisposing or protective alleles. The presence of a variable number and sequence polymorphism within the PATRR sequences in both LCR22-A and -B hint that this may be the case. In addition, with fully sequenced LCR22s it becomes possible to explore how the LCR22 repeat variability, orientation and rearrangements affect the 22q11.2DS phenotype.

## Materials and Methods

### Sample collection

A total of 49 Epstein-Barr virus transformed (EBV) cell lines, of which 25 were index patients with a *de novo* 22q11.2 deletion and 24 were parents-of-origin, were collected for the study (**Supplemental_Table_S1**). Four samples were collected from Albert Einstein College of Medicine (New York), 24 from University of Toronto, 2 from Children’s Hospital of Philadelphia and 21 from University Hospital Leuven . Seven of the LCR22-A/D deletion duos (AD009-AD015, **Supplemental_Table_S1**) were previously used in the study of Demaerel et al. (2019), and here their patterns were re-analyzed for crossover site delineation. All patients and parents had given written consent to participate in genetic research of the 22q11.2 CNV. The EBV cell lines were the starting material for the fiber-FISH and sequencing sample preparation. Study approval was obtained from the Medical Ethics Committee of the University Hospital/KU Leuven (S52418), and at the Institutional Review Boards of the originating sites of the participating families: Clinical Genetics Research Program at the Centre for Addiction and Mental Health (REB# 114/2001-02), the Albert Einstein College of Medicine (IRB# 1999-201-047), and Children’s Hospital of Philadelphia (IRB protocol #07-005352). Additional information regarding the samples is available in **Table S1**.

### Fiber-FISH & optical mapping

To haplotype the LCRs on chromosome 22, we used the LCR22-specific fiber-FISH method as described in Demaerel et al. (2019). In short, long DNA fibers were extracted from EBV cell lines from probands and the parent-of-origin using the Genomic Vision extraction kit (Genomic Vision). These long DNA molecules were combed onto slides and hybridized using a LCR22-specific customized probe set (Demaerel et al. 2019). Following automated microscopy scanning of the slides (FiberVision, Genomic Vision), the data were analyzed by manually indicating regions of interest (FiberStudio, Genomic Vision). Haplotypes were *de novo* assembled, with a coverage of at least 5X, using matching colors and distances between the probes as anchors. Patterns of recombined LCR22s of the patients were compared to the parental patterns to identify the haplotype alteration position. Optical mapping data was obtained for the AD004 Parent. DNA was extracted using the SP-G2 Blood & Cell culture DNA Isolation kit (#80060, Bionano Genomics) and labeled using the DLS-G2 DNA labeling Kit (DLE-1 labeling enzyme, #80046, Bionano Genomics). The sample was loaded onto Saphyr Chips G2.3 (Bionano Genomics, linearized and visualized using the Saphyr Instrument (Bionano Genomics), according to the System User Guide. The analysis was performed by *de novo* assembly against the hg38 reference genome in Bionano Access Software (Bionano Genomics).

### (Ultra-)long read sequencing via Oxford Nanopore Technologies (ONT)

Ultra-high molecular weight (UHMW) DNA (50kb -1Mb) was extracted via the UHMW DNA extraction protocol of the Nanobind CBB Big DNA kit (Circulomics) or via the Monarch HMW DNA Extraction Kit for Cells & Blood (New England Biolabs) and quantified using Qubit dsDNA Broad Range kit (ThermoFisher). Approximately 40µg was used as input for sequencing (SQK-ULK001 library preparation kit, ONT). The UHMW DNA was tagmented and adapters attached to the DNA ends, followed by a disk-based clean-up reaction or spermine precipitation (SQK-ULK001, ONT). One third of the library was loaded onto a Promethion flow cell (ONT). The flow cells were washed twice and reloaded with the remaining 2/3 of the library after 24h and 48h. Run statistics are presented in **Supplemental_Table_S2**.

### *De novo* assembly and sequence alignment

Ultra-long Nanopore reads were aligned to the human reference genome (hg38) with Minimap2 (Li 2018). To facilitate visualization of the alignment with with the Integrative Genomics Viewer (IGV, (Robinson et al. 2011), the 22q11 region was isolated with samtools (Li et al. 2009). *De novo* assembly was performed with NOVOLoci, a targeted haplotype-aware assembler (available at https://github.com/ndierckx/NOVOLoci, unpublished). NOVOLoci requires a seed sequence to initiate the assembly and produces separate assemblies for each haplotype. For the CTLR-Seq libraries, the target sequences served as seed sequences for the assemblies. For the whole-genome libraries, non-duplicated sequences downstream from the target regions were selected. To identify the rearrangement region, a multiple alignment between shared subunits among the two parental alleles was conducted using MAFFT (Katoh et al. 2019). A custom script was used to identify unique SNPs between these shared subunits, facilitating the identification of the transition between the two LCR22s.

### Deletion (and inversion) breakpoint identification in B2-B2 LCR22-A/B patterns

In patients and parents of families AB001, AB002, and AB004, reads (partly) covering the wild type and the rearranged LCR22s were manually selected based on LCR22 flanking sequence. In the parents-of-origin, two haplotypes for LCR22-A and -B could be differentiated, based on SNPs in the flanking sequence. The composition of the reads was determined using BLAT (Kent 2002), and the repeat composition between the two B2 probes in the rearranged allele was determined using RepeatMasker (Smit et al.).

### Long-range PCR over the LCR22-A/B recombination site and PacBio sequencing

Long-range PCR was performed using the TaKaRa LA PCR kit (TaKaRa Bio). PCR conditions were optimized for the extension-annealing phase, taking into account the presence of AT-repeats (Inagaki et al. 2005): aspecific bands are present when the temperature is below 60°C and there is no reaction above 63°C. A single primer (5’-ATACTACTGTGGCTTTGTTCCAAAG) was used as both forward and reverse primer. PCR was performed by an initial denaturation of 2 minutes at 94°C, 30 cycli of 30 seconds at 94°C followed by 7 minutes at 63°C, and the final elongation was at 60°C for 10 minutes. Fragments were analyzed on agarose gel.

A PacBio library was generated from the amplicons according to the Template Preparation and Sequencing protocol (Template Prep kit 3.0, Pacific Biosciences). Four libraries (22q11.2 patient AB004, his two children with a 22q11.2 deletion and the mother of the children) were pooled and loaded onto a single SMRT cells on a PacBio RSII using a DNA/polymerase binding kit P6 v2 (Pacific Biosciences) loading concentration 25pM) and DNA Sequencing Reagent kit 4.0 v2 (Pacific Biosciences). The RS_Long_Amplicon_Analysis.1 pipeline was used for analysis.

### Low-pass sequencing

The mosaic LCR22-A/B (AB002) deletion was detected in context of non-invasive prenatal testing of the pregnant patient JV2001. Procedures for non-invasive prenatal testing were followed as described (Bayindir et al. 2015).

### Array comparative genomic hybridization (ArrayCGH)

ArrayCGH was performed using the 60k CyoSure Constitutional v3 array (Oxford Gene Technology). Data analysis and visualization of the results was done using CytoSure Interpret Software (v4.10.44) with embedded Circular Binary Segmentation algorithm for automated copy number calling. The analysis was performed using hg19/GRCh37 genome build.

### SNP array

Genotyping was performed using Illumina HumanCytoSNP-12 BeadChip according to the Illumina Infinium HD Ultra protocol. Genotype, logR ratio and B-allele frequency (BAF) were extracted from the raw intensity data using the GenomeStudio software (v2.0.5) with the embedded genotype calling algorithm.

### Interphase FISH

Dual-color interphase-FISH was performed using Vysis DiGeorge LSI TUPLE1 (HIRA) Spectrum Orange / LSI ARSA Spectrum Green probe set (Abbott). 100 nuclei from blood, urine, and buccal mucosa were scored, by assessing the presence or absence of the Spectrum Orange fluorescent probe targeting the 22q11.2 HIRA region.

### Targeted interphase-FISH

BAC DNA was extracted from BAC clones (BacPac Resources, CHORI, Oakland) using the Nucleobond Xtra BAC kit (Macherey-Nagel) and subsequently labeled (Nick translation protocol, Abbott Molecular Inc.) (**Table 4**). EBV cells of the AB002 patient, a normal control, and a non-mosaic heterozygous LCR22-A/B deletion patient were fixed and slides for FISH prepared to score the inversion and deletion frequency. For family BD001, slides were prepared from the patient and both parents of the family. Aside from the blue (CH17-320A22) and/or orange (RP11-354K13) BAC probe that were used as control, the presence and orientation of the green (CH17-222C16) and red (CH17-389E17) BAC probe were essential to score a chromosome as normal, inversion, or deletion (**Supplemental_Table_S3**).

**Table 4:** Extracted BAC clones with LCR22-related location and labeling. The two last columns indicate whether the probe was used to score inversion status in the corresponding families.

| BAC clone | Location | Labeling | AB002 | BD001 |
| --- | --- | --- | --- | --- |
| CH17-320A22 | Proximal LCR22-A | Aqua 431dUTP | x | x |
| CH17-222C16 | Distal LCR22-A | Spectrum Green | x | x |
| CH17-389E17 | Proximal LCR22-B | Spectrum Orange | x | x |
| RP11-354K13 | Distal LCR22-D | Spectrum Green + Spectrum Orange | x |  |

### Etnicity determination

Data from 1000genomes phase3 were downloaded, converted to PLINK format and pruned to remove variants in linkage disequilibrium using PLINK (v1.9). The VCF files of the 22q11 samples have been generated by mapping the reads to reference genome (T2T) with minimap2 (v2.25) and variant calling with the PEPPER-Margin-DeepVariant pipeline (v0.8). The VCF files have then been lifted over to hg38 using Picard (v2.18.23), normalized and converted to BCF using BCFtools (v1.9), converted to PLINK format, merged and pruned to remove variants in linkage disequilibrium using PLINK (v1.9). Variants successfully genotyped in at least 90% of the 22q11 samples and available in 1000genomes phase3 dataset have been selected to perform a principal component analysis using PLINK (v1.9). A total of 646 variants could be used for the analysis. PCA analysis shows that the samples are of European descent, although American influences cannot be excluded (**Supplemental_Fig_S4**)

### Data access

Fiber FISH data, and cell lines used to map repeats are available upon request. Scripts used in this study are available at https://github.com/ndierckx/NOVOLoci

### Competing interest statement

The authors declare to have no competing interests.

## Acknowledgements

This work was supported by funds from the National Institute of General Medical Sciences (GM125757 to B.S.E).

## Author contributions

L.V. lead conception of the work and experimental design using fiber FISH and ONT. A.S., E.V informed parents and patients during hospital follow-up and collected blood samples. Data was collected by L.V., M.S.S., S.M., R.C., A.S., H.V.E., J.B. and L.V., N.D. and E.S. performed data analysis and interpretation. J.R.V., L.V., N.D., M.S.S., and E.S. drafted the article, which was critically revised by J.R.V., B.M., B.S.E., T.H.S., A.S., M.X., M.S.H., D.M.M. Final approval of the version to be published was given by L.V., N.D., M.S.S, S.M., E.S., R.C., T.H., K.D., H.P., D.M.M., A.S., J.B., B.S.E., H.V.E., A.S.B. and J.R.V.

## Notes

### Competing Interest Statement

The authors have declared no competing interest.

